# Age-related changes of deep-brain neurophysiological activity

**DOI:** 10.1101/2022.04.27.489652

**Authors:** T. Hinault, S. Baillet, SM. Courtney

## Abstract

Cognitive decline with age is associated with brain atrophy and reduced brain activations, but the underlying neurophysiological mechanisms are unclear, especially in deeper brain structures primarily affected by healthy aging or neurodegenerative processes. Here, we characterize time-resolved, resting-state magnetoencephalography activity of the hippocampus and subcortical brain regions in a large cohort of healthy young and older volunteers from the Cam-CAN open repository. The data show age-related changes in both rhythmic and arrhythmic signal strength and temporal variability in multiple deeper brain regions, including the hippocampus, striatum, and thalamus. We observe a slowing of neural activity in deeper brain regions, which echoes previous reports of cortical slowing. We also report reduced occipito-parietal alpha peak associated with increased theta-band activity and signal variability in the hippocampus, an effect that may reflect compensatory processes as theta activity and variability were more strongly expressed when cognitive performances are preserved. Overall, this study advances the understanding of the biological nature of inter-individual variability in aging. The data provide new insight of how hippocampus and subcortical neurophysiological activity evolve with biological age, and highlight frequency-specific effects associated with cognitive decline vs. cognitive maintenance.

## Introduction

The course of healthy aging is associated with preserved daily life autonomy, although the efficiency of cognitive functions such as memory and executive control diminishes (Hinault & Lemaire, 2020). Age-related cognitive decline is heterogeneous across individuals as brain functions are affected differentially across the population from the same age group (Reuter-Lorenz & Park, 2014). Cognitive decline is associated with brain atrophy and reduced task-related cortical activations relative to younger individuals (Spreng & Turner, 2019). However, the modifications of deep-brain neurophysiological activity with age are seldom investigated, despite the role of hippocampus and subcortical structures in cognitive functioning (Bourgeois et al., 2020), and their early involvement in neurodegenerative pathologies (Gulyaeva, 2019).

Task-free, spontaneous fluctuations of brain activity at rest have long been considered as unwanted background noise, yet recent works highlight the potential of neurophysiological signal dynamics as useful indicators of individual cognitive performance (Uddin, 2020; Waschke et al., 2021; Wiesman et al., 2021) and individual differentiation (da Silva Castanheira et al., 2021). The increased temporal variability of fMRI cortical and subcortical brain signals with biological age is negatively associated with cognitive performance (Guitart-Masip et al., 2016; Scarapicchia et al., 2019). Some of the rich dynamical features of neurophysiological activity accessible with magnetoencephalography and electroencephalography (M/EEG; Buzsáki, 2019, Baillet, 2017) change with biological age (Cheng et al., 2015; Courtney & Hinault, 2021). Indeed, the aging brain expresses increased slower activity below 4 Hz (delta frequency band), and increased temporal variability, both negatively associated with cognitive performance (Kumral et al., 2020; Jauny et al., 2022). Recent tools have emphasized the distinction between periodic (oscillatory) and aperiodic (background) signal components in electrophysiology (Donoghue et al., 2020; Voytek et al., 2015). The magnitude of aperiodic activity increases with aging (Merkin et al., 2021), a possible expression of increased neural noise associated with cognitive decline (Thuwal et al., 2021). However, most of these age-related electrophysiological observations so far are from neocortical activity.

Here, we sought to identify specific deep-brain neurophysiological signal features associated with the heterogeneity of cognitive aging. A growing body of work demonstrates MEG effects stemming from deeper structures (Coffey et al., 2016; Gorina-Careta et al., 2021; Müller et al., 2019; Samuelsson et al., 2021). We retrieved MEG data from younger and older individuals from the Cam-CAN (*Cambridge Centre for Ageing and Neuroscience*) repository to investigate time-resolved resting state neurophysiological fluctuations in the hippocampus, striatum and thalamus.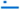Because these structures are associated with age-related alterations of short-term and working memory functions (O’Shea et al., 2016; Valdés Hernández et al., 2020), we expected that they expressed slower, reduced or more variable neurophysiological activity in older adults._We also hypothesized that preserved cognitive performances in healthy older adults, would also be associated with increased deep-brain activity.

## Results

### Behavioral differences between age groups

The younger and older adults’ groups had similar biological sex ratios and showed normal general cognitive performance (above the MMSE > 27/30 cutoff; see Table 1).

**Table 1.** Demographic and cognitive characteristics [average (standard deviation)] of the study participants retrieved from the Cam-CAN dataset.

| Variables | Young adults | Older adults | *F* | *p* |
| --- | --- | --- | --- | --- |
| *N* | 47 | 47 | - | - |
| Females/males | 30/17 | 30/17 | - | - |
| Age in years | 26 (2.0) | 73 (2.7) | - | - |
| Years of Education | 16 (2.8) | 13 (4.5) | 12.45 | 0.001 |
| Mini Mental State Evaluation (MMSE) | 29.51 (0.9) | 28.43 (1.2) | 25.65 | <0.001 |
| Visual short-term memory (accuracy) | 0.49 (0.1) | 0.42 (0.1) | 21.19 | <0.001 |
| Individual Alpha Frequency (IAF, in Hz) | 10.10 (0.9) | 9.21 (0.7) | 24.73 | <0.001 |

### Between-group differences in neurophysiological activity

IAF, recorded over parieto-occipital sensors, was lower in older adults relative to younger adults (Table 1). We found steeper slopes of arrhythmic activity in older adults relative to younger adults, in all ROIs. That is, delta-band activity was larger, with reduced gamma-band activity, in older adults (Figure 1A-B), in line with the slowing of the dominant activity reported at the sensor/cortical level. A similar pattern was observed regarding the signal temporal variability (Figure 1C-D). Larger offsets were also observed in bilateral striatum and hippocampus (Figure 2A-B). Importantly, in the right hippocampus only, theta-band activity and variability were stronger in older adults. Beta-band activity was also reduced in older adults, bilaterally in nucleus accumbens, and in the right putamen. Control analyses replicated previously reported effects of cortical increases of delta-band activity and reduced gamma-band activity at rest across cortical regions. Increased theta-band activity and variability, however, were not significant in cortical regions, suggesting that the changes observed in the hippocampus are specific to that structure.

**Figure 1.**
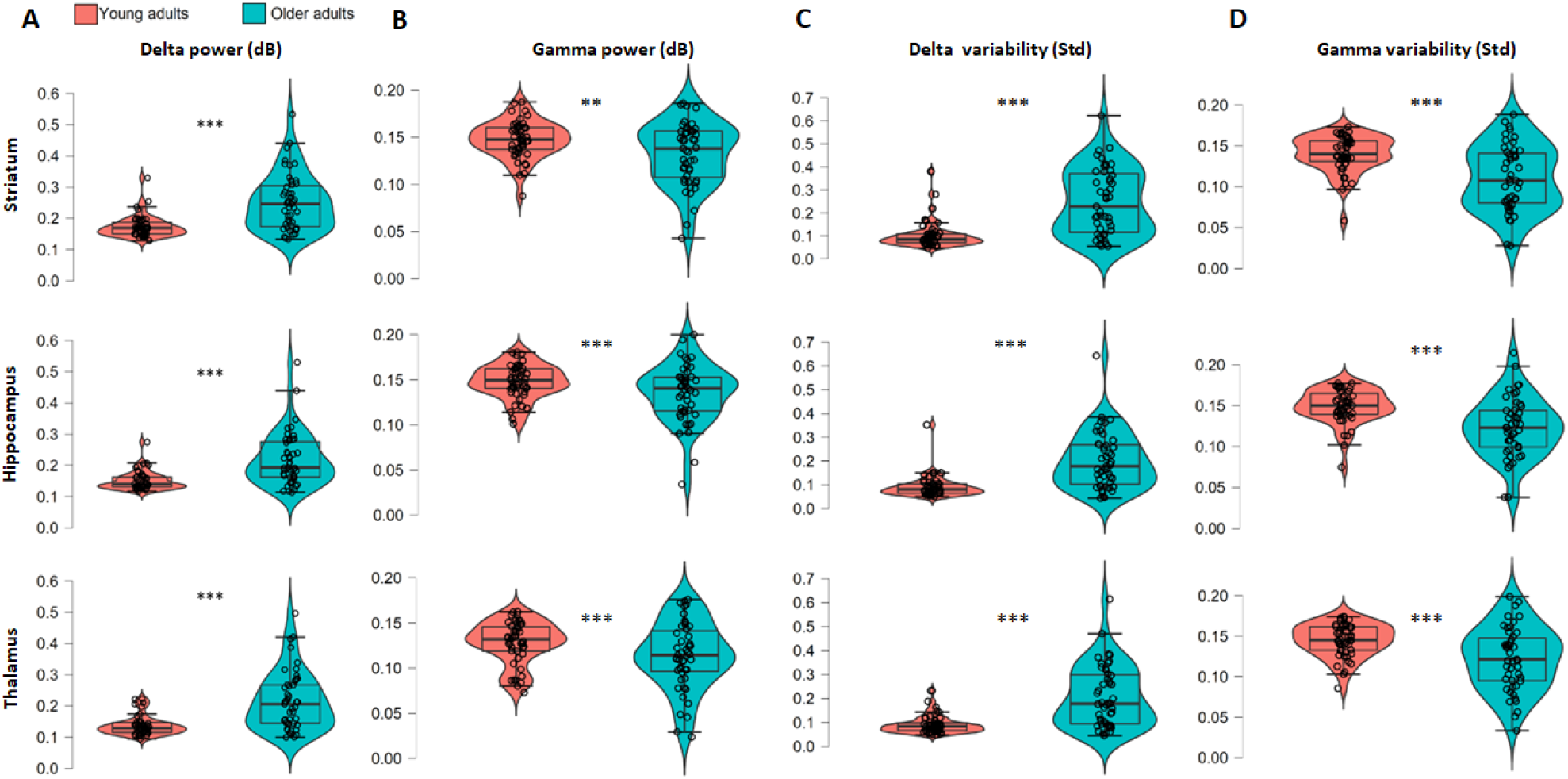
Bilaterally in all tested ROIs of older adults relative to younger adults: A) Delta-band activity was stronger; B) Gamma-band activity was reduced; C) Delta-band temporal variability was larger; D) Gamma-band temporal variability was smaller. The plots show the average regional activity between left and right homologous structure

**Figure 2.**
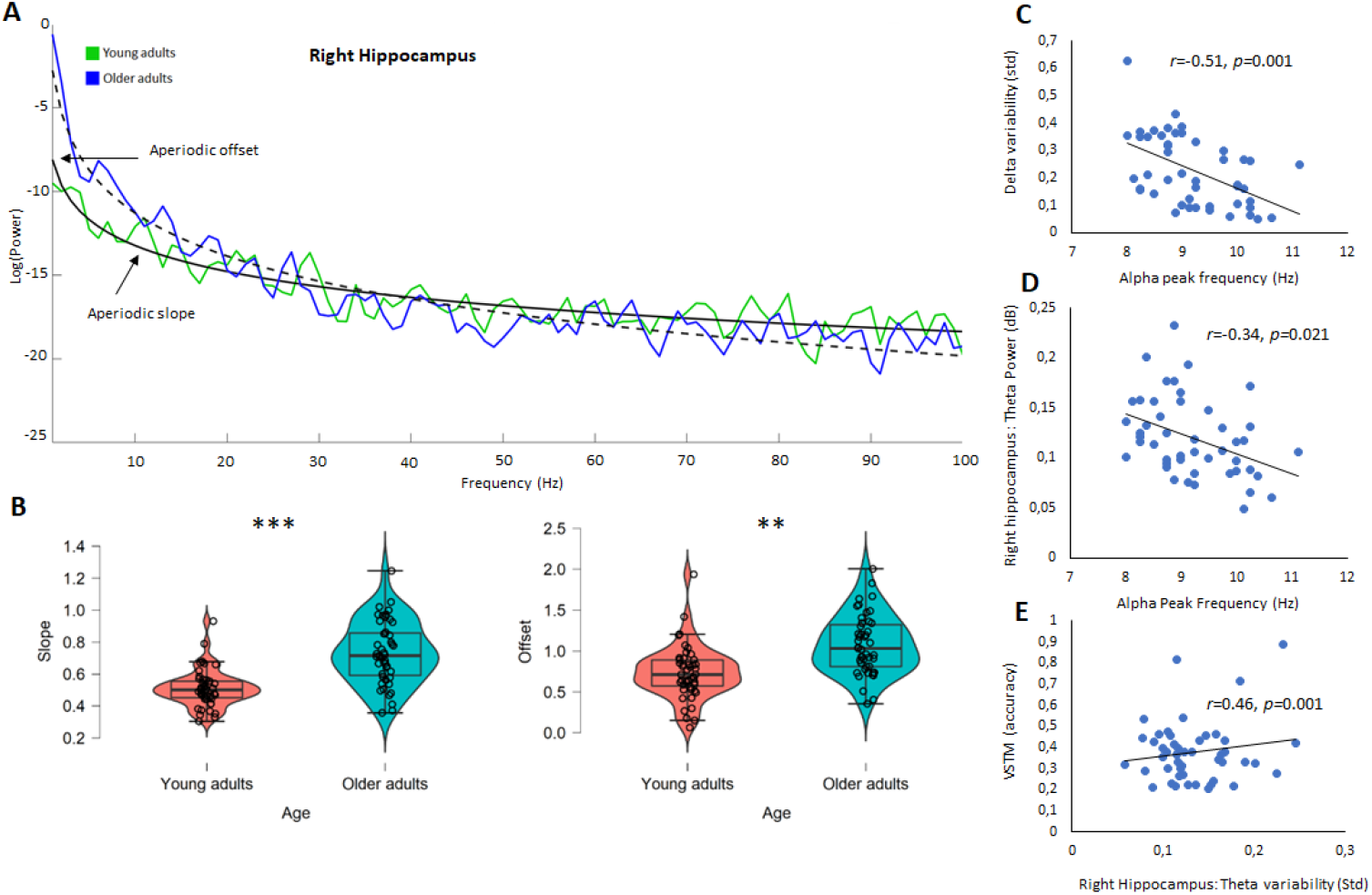
A) Parametrization of the group average power spectral density in young and older adults in the right hippocampus region, showing the aperiodic slope and offset across the frequency range. B) Steeper slopes (p<0.001) and larger offsets (p=0.007) of the aperiodic spectral component were observed in older adults relatively to young adults, from resting-state activity of the right hippocampus. C) Negative association between the individual occipito-parietal alpha peak frequency and deep-brain delta variability (averaged across regions) in older adults. D) Negative association between the individual alpha peak frequency and power of theta-band neurophysiological activity of the right hippocampus in older adults. E) Positive association between theta variability of neurophysiological activity in the right hippocampus and VSTM task performances of older adults.

No correlation between neurophysiological activity measures was observed in young adults. In older adults, a lower occipito-parietal IAF was associated with stronger deep-brain delta-band activity (all *p*<0.009), and temporal variability in all ROIs (all *p*<0.005; Figure 2C-D). Importantly, a lower surface IAF was also associated with a stronger right hippocampus theta activity (*r*=-34, *p*=0.021).

### Associations of deep-brain neurophysiological activity with cognition in older adults

Stronger right hippocampus theta activity was associated with higher VSTM performance in older adults (*r*=0.49, *p*<0.001, respectively). VSTM performance was also positively associated with temporal variability in the theta band in the right hippocampus and left thalamus (*r*=0,46, *p*=0.001, and *r*=0,52, *p*<0.001, respectively). We also found a positive association between the slopes of arrhythmic activity in the right hippocampus and VSTM performances (*r*=0.62, *p*<0.001; Figure 2E).

## Discussion

We believe this study is first to report aging effects on human deep-brain neurophysiological activity and signal variability over time. We used recent methodological advances in source modelling of resting-state M/EEG signals (Samuelsson et al., 2021), and spectral parametrization of periodic and aperiodic brain neurophysiological activity (Donoghue et al., 2020). Overall, older adults showed stronger and more temporally-variable neurophysiological activity in the alpha and lower frequency bands (Figure 3). These effects were reversed in higher frequency bands. Most age-related changes reported in the present study are associated with cognitive decline; however, we also found that theta-band signal strength and temporal variability in the right hippocampus were positively associated with VSTM performance. Taken together, our data show that biological aging does impact subcortical and hippocampus neurophysiological activity, with differential consequences on cognitive performances.

**Figure 3.**
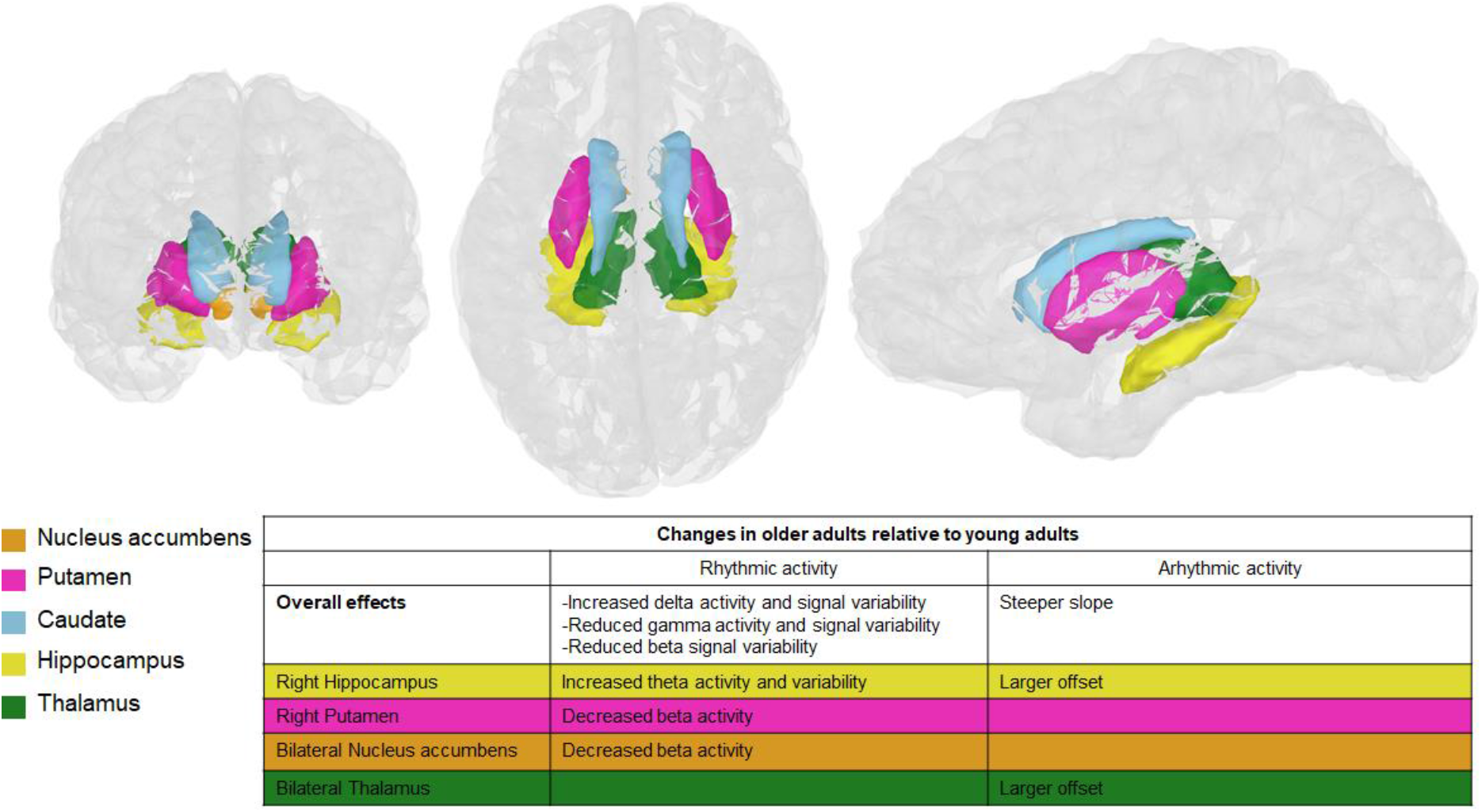
Summary of the reported frequency-specific and aperiodic effects found in older adults relative to young adults, across deep-brain regions of interest.

Our study replicates and extends previous findings of decreased IAF measured from scalp (Scally et al., 2018), and of cortical slowing across the typical frequency bands of electrophysiology with age (e.g., Courtney & Hinault, 2021). Our present results point at possible deeper brain origins of such overall slowing of brain activity, and are in line with fMRI findings of age-related subcortical changes (Daugherty et al., 2015). Our data show that with aging, subcortical signal variability is reduced in older adults for the fastest frequency bands. Symmetrically, temporal variability was more pronounced in the slower delta frequency band, in association with reduced IAF. These effects were associated with decreased cognitive performances.

Our study disentangles between age-related effects on periodic vs. aperiodic deeper brain activity. Changes in aperiodic cortical signal power have been reported (Merkin et al., 2021; Thuwal et al., 2021; Voytek et al., 2015), and discussed as reflecting larger amounts of neural noise in brain communications with advancing age. Age-related brain noise is possibly related to alterations of the excitation/inhibition balance in neural circuits (Donoghue et al., 2020; Voytek & Knight, 2015), and would impact information processing and the efficiency of cognitive processes (Merkin et al., 2021). Our data specifies how age affects the spectral slope and offset of aperiodic neurophysiological activity in the hippocampus and subcortical regions.

The hippocampus generates theta activity (e.g., Goutagny et al., 2009), in association with memory processes such as, encoding (Fell et al., 2011), short-term and working memory (Axmacher et al., 2010). Here, we found that both the spectral slope and theta-band temporal variability of hippocampus activity are positively associated with VSTM performance. In a similar fashion, we found that increased theta-band signal variability in the thalamus was also positively associated with VSTM performance.

Our data show that preserved cognitive performance with advancing age is associated with both the strength and variability of hippocampus theta-band rhythmic activity. This aspect is in line with previous fMRI work that showed an association between preserved cognitive performance in older adults and higher hippocampal activity (e.g., Lister & Barnes, 2009). Our data show that preserved cognitive performance with advancing age is associated with both the strength and variability of hippocampus theta-band rhythmic activity. These theta-band changes were not observed in control cortical regions, suggesting that these changes (but not changes in other frequency bands) are hippocampal-specific. We anticipate that these results contribute to future investigations of the inter-individual variability in cognitive performances in aging (Cabeza et al., 2018). We emphasize though that hyper-activity is not always of a compensatory nature (Hillary & Grafman, 2017) and may be indicative of subsequent cognitive decline. Future longitudinal studies will need to clarify these aspects.

We also discuss some identified limitations to the present study. Courtney and Hinault (2021) and Finn (2021) recently discussed how resting-state activity is less directly associated with cognitive functioning than task-related activity. Moreover, the Cam-CAN cognitive tests did not specifically target executive functions that are associated with frequency-specific brain activity (Hinault et al., 2020, 2021). Finally, the investigation of age-related changes in deep-brain activity would benefit from the specification of longitudinal trajectories of brain changes, which was not possible with the present dataset.

Recent fMRI work has highlighted hippocampus and subcortical brain activity in higher-order cognitive functions (Bourgeois et al., 2020; Chiu & Egner, 2019). Our results reveal that aging effects on these regions are characterized by changes in both rhythmic and arrhythmic signal strength and temporal variability, with an overall slowing of deeper neural activity. Individual differences were also observed, with specific increased theta-band activity and variability in the hippocampus associated with preserved short-term memory performances. Relative to healthy aging, the deep-brain regions investigated here are further impaired in age-related pathologies (Gulyaeva, 2019), which we anticipate may be associated with further alterations of their neurophysiological activity.

## Methods

### Participants’ characteristics

The structural MRI and resting-state MEG data from the Cam-CAN repository (Shafto et al., 2014; Taylor et al., 2017) were available from http://www.mrc-cbu.cam.ac.uk/datasets/camcan/). The retrieved sample consisted of forty-seven young adults (20-30 years, 30 females) and forty-seven older adults (65-75 years, 30 females; Table 1). Details on the demographic and behavioral data are available online (https://camcan-archive.mrc-cbu.cam.ac.uk/dataaccess/). All older adults scored within normal range at the Mini Mental State Evaluation (MMSE score > 27; Folstein et al., 1975). No participants reported a history of neurological or cognitive disorders, traumatic brain injury, or major psychiatric disorders.

### Behavioral tasks

Because deep-brain regions are essential to short-term memory and working memory performance (McNab & Klingberg, 2008), we investigated age-related changes in the visual short-term memory (VSTM) task. In this task, participants were briefly presented 1 to 4 colored discs on a computer screen and asked to recall the color of the target disc at a cued location. Participants reported their delayed response on a color wheel using a touchscreen input (further detail concerning the task is available online;https://camcan-archive.mrc-cbu.cam.ac.uk/dataaccess/).

### Neuroimaging data

The MRI data consisted of T1-weighted image volumes (field of view: 256×240×192 mm, 1×1×1 mm voxel size, repetition time: 2250 ms, echo time: 900 ms, flip angle: 9°). MEG data consisted of approximately 9 minutes of eyes-closed resting-state recordings acquired with a 306-channel Elekta Neuromag Vectorview MEG system (102 magnetometers and 204 planar gradiometers), with 1kHz sampling rate, and a 0.03-330 Hz online bandpass filter. Registration of MEG and MRI used digitized anatomical landmarks (i.e., fiducial points; nasion and left/right preauricular points, and additional scalp points). The electro-occulogram (EOG) and electrocardiogram (ECG) were recorded to capture eye movements and heartbeats, respectively.

### MEG data analysis

The retrieved data were already partly preprocessed using the temporal signal space separation approach (tSSS): 0.98 correlation, 10 s window; bad channel correction: ON; motion correction: OFF; 50Hz+harmonics (mains) notch. We performed further artifact detection and attenuation (on continuous data), filtering (0.3-100 Hz bandpass, also on continuous data), and source estimation using *Brainstorm* (Tadel et al., 2011), all with default parameters, unless noted below. Remaining physiological artifacts (e.g., eye blinks and saccades) were identified and removed with bespoke signal-space projections (Uusitalo & Ilmoniemi, 1997). For each participant, we identified defective sensors and individual alpha peak frequencies (IAF) using power-spectrum density (PSD) estimates (Welch’s method) from their entire recording. We defined IAF as the average frequency of the PSD spectral peaks in the alpha frequency range (8-13 Hz) observed over occipito-parietal gradiometers and magnetometers. To account for individual brain atrophy and inform MEG source imaging (Baillet, 2017), we used *Freesurfer* (Fischl, 2012) to produce segmentations of head tissues, including cortex and subcortical structures, and to compute the intracranial volume from each participant’s MRI.

We modeled MEG brain source activity using elementary volume current dipoles (15,000 elementary source locations across the entire brain without orientation constraint; Baillet et al., 2001). To that end, we used the Desikan/Kiliany atlas (*aparc+aseg* segmentation; Desikan et al., 2006) of *Freesurfer’s* individual cortical and subcortical parcellations. We then used the overlapping-sphere approach to MEG forward head modelling (Baillet et al., 2001). We could not explicitly account for environmental and instrumental noise, as empty-room recordings were not available in Cam-CAN. We derived MEG imaging kernels for each individual using the dynamic Statistical Parametric Mapping approach (dSPM; Dale et al., 2000; Hauk et al., 2011) to estimate the source time series of each region of interest (ROI; left and right hippocampus, thalamus, nucleus accumbens, caudate, and putamen). To determine the specificity of deep-brain effects, we performed control analyses on a subset of cortical regions of the *aparc* atlas, which were previously reported as showing changes in neurophysiological activity associated with age-related short-term and working memory performance alterations (e.g., Hinault et al., 2020).

We performed time-frequency decompositions of MEG source time series using the Hilbert transform in frequency bands of interest. The width of each frequency band was based on the surface IAF value of each participant (Toppi et al., 2018): delta (IAF-8/IAF-6), theta (IAF-6/IAF-2), alpha (IAF-2/IAF+2), beta (IAF+2/IAF+14), low-gamma (IAF+15/IAF+30), and high-gamma (IAF+31/IAF+90). We used the first principal component of the time series within each subcortical ROI as a summary statistic of distributed neurophysiological activity, which reduces cross-talk between regions (Sato et al., 2018). Following the Hilbert transform, signals were normalized for group comparisons through spectrum normalization. To determine whether the temporal variability of resting-state activity was affected by physiological aging, we considered the standard deviation of time series (Coquelet et al., 2017) of deep-brain neurophysiological activity.

We used *specparam* to parametrize the time-frequency signal of each ROI (Donoghue et al., 2020) and measure the offset and slope model parameters of aperiodic spectral components (Merkin et al., 2021). Offset reflects a shift of signal power across frequencies, and slope accounts for the steepness of the decrease of broadband background signal power with frequency.

We used non-parametric inferential statistics based on permutation tests, false discovery rate (FDR) corrected (N=10,000; Maris & Oostenveld, 2007) and correlation analyses, also FDR-corrected. To limit the number of comparisons, only regions and frequencies showing significant between-group differences in permutation tests (p<0.05, FDR correction over signal, time and frequency dimensions) were further considered for possible association with cognitive performance.

## Acknowledgments

We would like to thank Gwendolyn Jauny for her help with data preprocessing. Data collection and sharing for this project were partly provided by the Cambridge Centre for Ageing and Neuroscience (Cam-CAN). Cam-CAN funding was provided by the UK Biotechnology and Biological Sciences Research Council (grant number BB/H008217/1), together with support from the UK Medical Research Council and University of Cambridge, UK. S.B. is supported by a NSERC Discovery grant, the Healthy Brains for Healthy Lives initiative of McGill University under the Canada First Research Excellence Fund, and the CIHR Canada Research Chair of Neural Dynamics of Brain Systems.

## Author contribution

Thomas Hinault designed the research, collected, and analyzed the data, and wrote the paper. Sylvain Baillet and Susan Courtney helped analyze the data and write the paper.

